# The predictive power of mRNA mapping for Cannabinoid 1 receptor protein in the human brain

**DOI:** 10.1101/2022.03.03.482632

**Authors:** Kyoungjune Pak, Tatu Kantonen, Laura Pekkarinen, Pirjo Nuutila, Lauri Nummenmaa

## Abstract

Type 1 cannabinoid (CB1) receptor is expressed in cortex, hippocampus, amygdala, basal ganglia, and cerebellum. With the help of the Allen Human Brain Atlas, genomic maps visualize not only the gene expression across whole brain regions, but also the functional profile of brain structures. Therefore, it is more timely than ever to integrate genomic mapping from brain mRNA atlas with the protein expression from positron emission tomography (PET) scans for better understanding of CB1 receptor of the human brain. F18-FMPEP-d2 PET scans were retrieved from the AIVO neuroinformatics project. Autoradiography data were based on the study with H3-CP55940. mRNA expressions of CNR1 gene (Cannabinoid receptor 1) were downloaded from the Allen Human Brain Atlas. Volume of distribution (V_T_) from F18-FMPEP-d2 PET scans, CNR1 gene expression, and H3-CP55940 binding were calculated and Spearman correlation analysis was performed. Also, a meta-analysis was done to investigate the association between protein expression from PET and mRNA expression from the Allen Human Brain Atlas. Between V_T_ of F18-FMPEP-d2 PET scans and CNR1 mRNA expression, moderate strength of correlation was observed *(rho* = 0.5026, p = 0.0354). Strong positive correlation was also found between CNR1 mRNA expression and H3-CP55940 binding (*rho* = 0.6727, p = 0.0281), validating the finding between F18-FMPEP-d2 PET scans and CNR1 mRNA. From the meta-analysis, the correlation coefficient ranged from −0.46 to 0.99, with a pooled effect of 0.58. In conclusion, the moderate to strong associations between gene and protein expression for CB1 receptor in the human brain were observed that CNR1 mRNA mapping might have the predictive power for in vivo CB1 receptor protein expression. From the meta-analysis, the moderate to strong correlation was observed between mRNA expression and protein expressions across multiple genes, showing the predictive power of genes to estimate protein levels of human brains.

**HIGHLIGHTS:**

- We investigated the association between CNR1 gene expression from the Allen Human Brain Atlas and type 1 cannabinoid (CB1) receptor expression from F18-FMPEP-d2 PET scans.
- The moderate to strong associations between gene and protein expression for CB1 receptor in the human brain were observed that CNR1 mRNA mapping might have the predictive power for in vivo CB1 receptor protein expression.
- From the meta-analysis, the moderate to strong correlation was observed between mRNA expression and protein expressions across multiple genes, showing the predictive power of genes to estimate protein levels of human brains.

## 1. INTRODUCTION

The use of cannabinoid is of growing interest, especially after the legalization of the recreational use of cannabinoids in Canada from 2018 (Haines-Saah and Fischer, 2021). The medical use of cannabinoids is helpful in certain forms of epilepsy, chemotherapy related vomiting, chronic pain, however, is still controversial over the medical use of cannabinoids due to inconsistent results and lack of evidence (Montero-Oleas et al., 2020). The endocannabinoid system consists of type 1 (CB1) and type 2 (CB2) cannabinoid receptors, endogenous ligands and their metabolic enzymes (Tao et al., 2020). CB1 receptor, encoded by CNR1 gene, is expressed in cortex, hippocampus, amygdala, basal ganglia, and cerebellum (Mackie, 2005). CB1 receptors are found primarily in the presynapses of the neurons (Mechoulam and Parker, 2013). They act as a neuromodulators, inhibiting the release of GABA, glutamate, or dopaminergic neurotransmitters into the synapse (Dickens et al., 2020; Piomelli, 2003). Alteration of CB1 receptor has reported in neuropsychological disorders, higher CB1 receptor availability in post-traumatic stress disorder, and lower CB1 receptor availability in schizophrenia (Neumeister et al., 2013; Ranganathan et al., 2016). Also, CB1 receptor antagonist, rimonabant was introduced as an anti-obesity drug, however, was withdrawn due to serious side effects including depressive disorders or mood alterations (Di Marzo and Despres, 2009). Therefore, CB1 receptor has been a target for drug development and in vivo imaging biomarker for neuropsychiatric disorders (Van Laere, 2007).

The Allen Human Brain Atlas is a freely available multimodal atlas of gene expression with visualization and data-mining resources that comprises a comprehensive array-based dataset of gene expression (Shen et al., 2012). With the help of the Allen Human Brain Atlas, genomic maps visualize not only the gene expression across whole brain regions, but also the functional profile of brain structures (Sandberg et al., 2000). Previous studies have proved the predictive power of brain mRNA mapping for in vivo protein expression from positron emission tomography (PET) for serotonin receptors (Beliveau et al., 2017; Komorowski et al., 2017; Rizzo et al., 2014), opioid receptors (Rizzo et al., 2014) and monoamine oxidase A (Komorowski et al., 2017; Zanotti-Fregonara et al., 2014), based on the key assumption that mRNA expression predicts protein expression. However, the studies on serotonin transporter or dopamine transporter showed a weak association between mRNA and protein levels, emphasizing the role of translational and post-translational mechanisms (Beliveau et al., 2017; Pak et al., 2022). To date there have been no published data on the association of CB1 gene expression with CB1 protein expression in humans. Previous studies of CB1 receptor has focused mainly on receptor binding, signal transduction, with lack of knowledge on CB1 gene regulation (Laprairie et al., 2012). Pharmacological exposures such as methamphetamine, alcohols, cannabinoids as well as physiological processes are known to modulate CB1 receptor mRNA expression (Laprairie et al., 2012). Therefore, it is more timely than ever to integrate genomic mapping from brain mRNA atlas with the protein expression from PET scans for better understanding of CB1 receptor of the human brain (Gryglewski et al., 2018). Here, we investigated the association between CNR1 gene expression from the Allen Human Brain Atlas and CB1 receptor expression from F18-FMPEP-d2 PET scans. In addition to protein expressions from PET scans, H3-CP55940 bindings from autoradiography (Glass et al., 1997) were collected, and analyzed to validate the findings of this study. Finally, to compare the predictive power of CB1 receptor with those of other neuroreceptor/transporters, we conducted a meta-analysis on the association between mRNA expression from the Allen Human Brain Atlas and protein expressions from multi-tracer scans.

## 2. MATERIALS AND METHODS

### 2.1. PET data acquisition

The study subjects were retrieved from the AIVO neuroinformatics project (http://aivo.utu.fi), in vivo molecular brain scan database hosted by Turku PET Centre. We identified all F18-FMPEP-d2 baseline PET studies of subjects without neurologic and psychiatric disorders, current use of medications that could affect CNS or abuse of alcohol or illicit drugs. Final sample consisted of 36 subjects of nonsmoking males. All F18-FMPEP-d2 scans were acquired with GE Discovery VCT PET/CT (Computed tomography) (GE Healthcare). The tracer was administered in a catheter placed in subject’s antecubital vein. CT scans were acquired before PET scans for attenuation correction. MR images (TR, 8.1 ms; TE, 3.7 ms; flip angle, 7°; scan time, 263 s; 1 mm3 isotropic voxels) were obtained with PET/MR (Ingenuity TF PET/MR, Philips) for anatomical normalization. The study was conducted in accordance with the Declaration of Helsinki and approved by the Turku University Hospital Clinical Research Services. The participants in this study were included in previous studies on feeding behavior and obesity (Kantonen et al., 2021a; Kantonen et al., 2021b). PET images were processed with automated processing tool Magia (https://github.com/tkkarjal/magia) (Karjalainen et al., 2020). Magia uses FreeSurfer (http://surfer.nmr.mgh.harvard.edu/) to define the regions of interest (ROIs). As there exists no suitable central reference region for F18-FMPEP-d2, volume of distribution (V_T_) (Terry et al., 2010), FMPEP-d2 V_T_ was quantified using graphical analysis by Logan (Logan, 2000). The frames starting 36 min and more after injection were used in the model fitting, since Logan plots became linear after 36 min (Logan, 2000). Plasma activities were corrected for plasma metabolites as described previously (Lahesmaa et al., 2018). From F18-FMPEP-d2 PET scans, V_T_ from 18 ROIs were extracted; amygdala, hippocampus, caudate, putamen, nucleus accumbens, thalamus, cerebellum, anterior cingulate cortex, posterior cingulate cortex, insula, orbitofrontal cortex, superior temporal cortex, mid temporal cortex, inferior temporal cortex, superior frontal cortex, midbrain, pons, medulla.

### 2.2. Autoradiography

Autoradiography data were based on the study by Glass et al. with H3-CP55940 (Glass et al., 1997). The data of H3-CP55940 binding from 8 adult human brains with a mean age of 55.6 years (range, 21-81 years) were averaged across layers for each ROI and analyzed with mRNA expression of CNR1 gene. From the results of autoradiography, H3-CP55940 binding was extracted from 11 ROIs; amygdala, hippocampus, caudate, putamen, nucleus accumbens, thalamus, cerebellum, mid temporal cortex, midbrain, pons, medulla.

### 2.3. mRNA data

The gene expression information investigated in this study was obtained from the freely available Allen Human Brain Atlas (www.brain-map.org) (Hawrylycz et al., 2012). We downloaded mRNA expressions of CNR1 gene (Cannabinoid receptor 1) to test for an association with the protein distribution. The dataset of the Allen Human Brain Atlas contains the gene expression profiles of CNR1 throughout the brain obtained from six healthy donors of 89 probes with a mean age of 42.5 years (range, 24-57 years). We downloaded mRNA expression of CNR1 gene in log2-values for each sample. For ROI-based analysis, mRNA expression values of CNR1 gene were obtained from the average within each region, and the median across probes.

### 2.4. Meta-analysis

We performed a systematic search of MEDLINE (from inception to November 2021) for articles published in English using the keywords “positron emission tomography,” “single photon emission computed tomography”, “mRNA,” and “Allen human brain atlas.” All searches were limited to human studies. The inclusion criteria were primary studies that analyzed the association between protein expression from PET/single photon emission computed tomography (SPECT) and mRNA expression from the Allen Human Brain Atlas. If there was more than one published study using the same dataset of PET scans, only one report with the information most relevant to this study was included. Two authors performed the searches, screening independently, and discrepancies were resolved by consensus. Data were extracted from the publications independently by two reviewers, and the following information was recorded: first author, year of publication, ROIs, the number of patients, criteria for inclusion/exclusion, mRNA, and radiopharmaceuticals. Effect size was the correlation coefficient with 95% confidence interval (CI). The electronic search identified 164 articles from MEDLINE. After excluding conference abstracts, animal studies, non-English studies, and studies that did not meet the inclusion criteria after screening the title or abstract, 11 studies were eligible for inclusion in this study (Beliveau et al., 2017; Kim et al., 2020; Komorowski et al., 2017; Komorowski et al., 2020; Lohith et al., 2017; Norgaard et al., 2021; Pak et al., 2022; Rizzo et al., 2016; Rizzo et al., 2014; Veronese et al., 2016; Zanotti-Fregonara et al., 2014).

### 2.5. Statistical analysis

Normality was assessed using the Shapiro-Wilks test. Autocorrelation of V_T_ from F18-FMPEP-d2 PET scans and CNR1 gene expression was assessed with Spearman correlation. For each ROI, mean V_T_ from F18-FMPEP-d2 PET scans, CNR1 gene expression, and H3-CP55940 binding was calculated and Spearman correlation analysis was performed to assess the association between them. All analyses were conducted using R Statistical Software (version R 4.1.1, The R Foundation for Statistical Computing).

## 3. RESULTS

Mean distribution of F18-FMPEP-d2 PET scans from 36 subjects are visualized in figure 1. The distribution of CNR1 mRNA expression and V_T_ of F18-FMPEP-d2 PET from each ROI are shown in figure 2. Strong auto-correlation was observed both for V_T_ from F18-FMPEP-d2 PET scans (inter-subject: mean correlation coefficient *rho*: 0.8765) and for CNR1 mRNA expression from the Allen Human Brain Atlas (inter-probe: mean correlation coefficient *rho* = 0.9138), which ensures consistency of observations. Between V_T_ of F18-FMPEP-d2 PET scans and CNR1 mRNA expression from 18 ROIs, moderate strength of correlation was observed (*rho* = 0.5026, p = 0.0354). Strong positive correlation was also found between CNR1 mRNA expression and H3-CP55940 binding from 11 ROIs (*rho* = 0.6727, p = 0.0281) (Figure 3), which validates the finding between F18-FMPEP-d2 PET scans and CNR1 mRNA.

**Figure 1.**
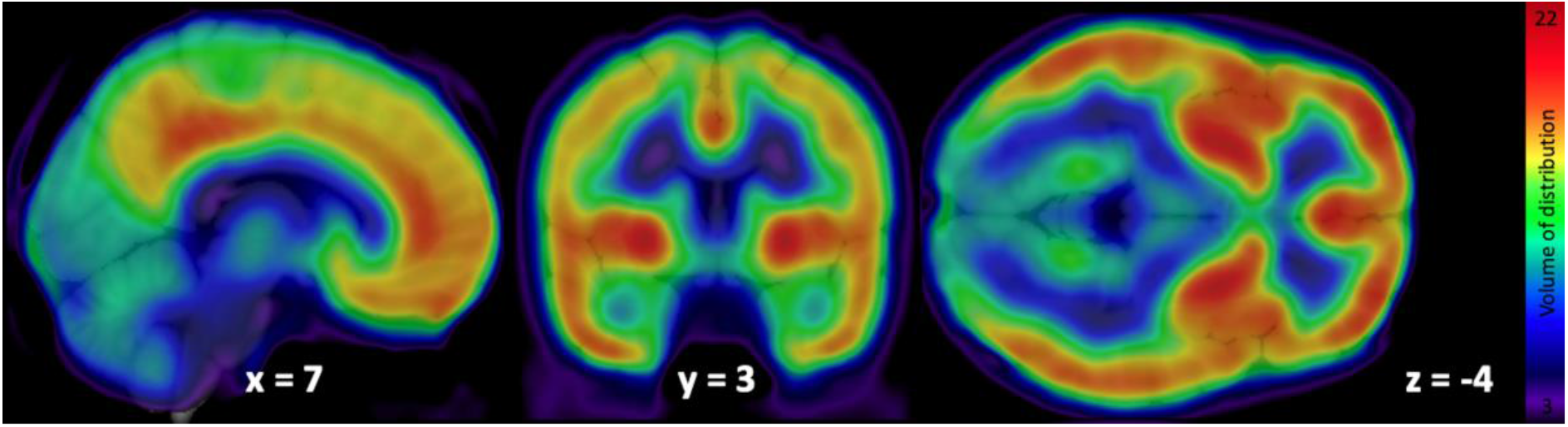
Mean volume of distribution of F18-FMPEP-d2 PET scans from 36 subjects

**Figure 2.**
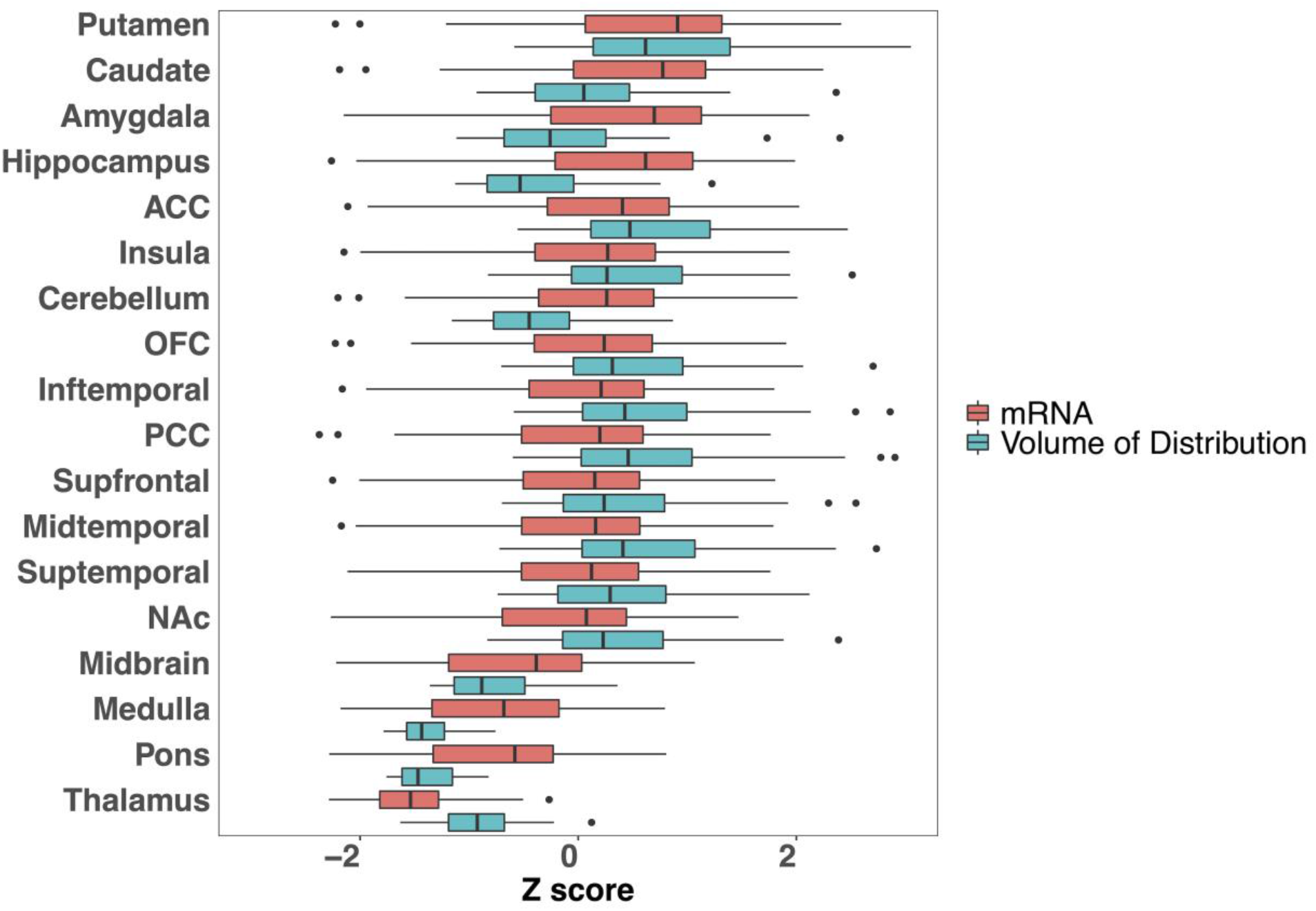
The distribution of CNR1 mRNA expression and volume of distribution of F18-FMPEP-d2 PET from 18 regions of interest. ACC, anterior cingulate cortex; OFC, orbitofrontal cortex; PCC, posterior cingulate cortex; NAc, nucleus accumbens.

**Figure 3.**
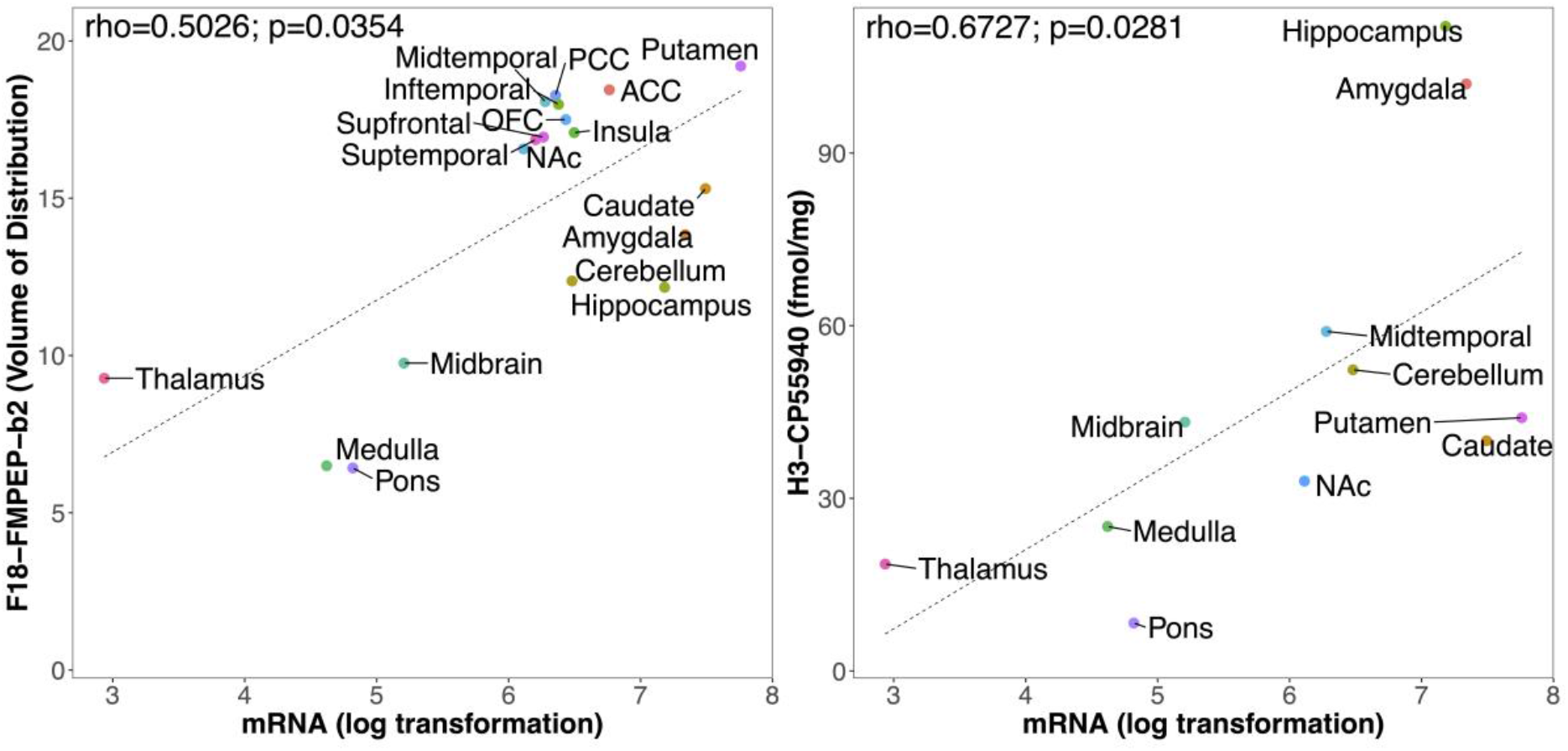
The association between CNR1 gene expression from the Allen Human Brain Atlas and CB1 receptor expression. Left: Between volume of distribution of F18-FMPEP-d2 PET scans and CNR1 mRNA expression from 18 regions of interest, moderate strength of correlation was observed *(rho* = 0.5026, p = 0.0354). Right: Between H3-CP55940 binding of autoradiography and CNR1 mRNA expression from 11 regions of interest, strong positive correlation was observed (*rho* = 0.6727, p = 0.0281).

The characteristics of studies included in the meta-analysis are summarized in table 1. The correlation coefficient ranged from −0.46 between DRD2 mRNA and C11-PHNO PET to 0.99 between GRM1 mRNA and F18-FIMX PET, with a pooled effect of 0.58 (95% CI: 0.47 ~ 0.69, I^2^=96.9%) (Figure 4). In a subgroup analysis, the pooled correlation coefficients were 0.41 (0 ~ 0.82, I^2^=91.8%) for dopaminergic system, and 0.76 (0.62 ~ 0.90, I^2^=86.0%) for serotonergic system, without a significant difference (p=0.1161).

**Figure 4.**
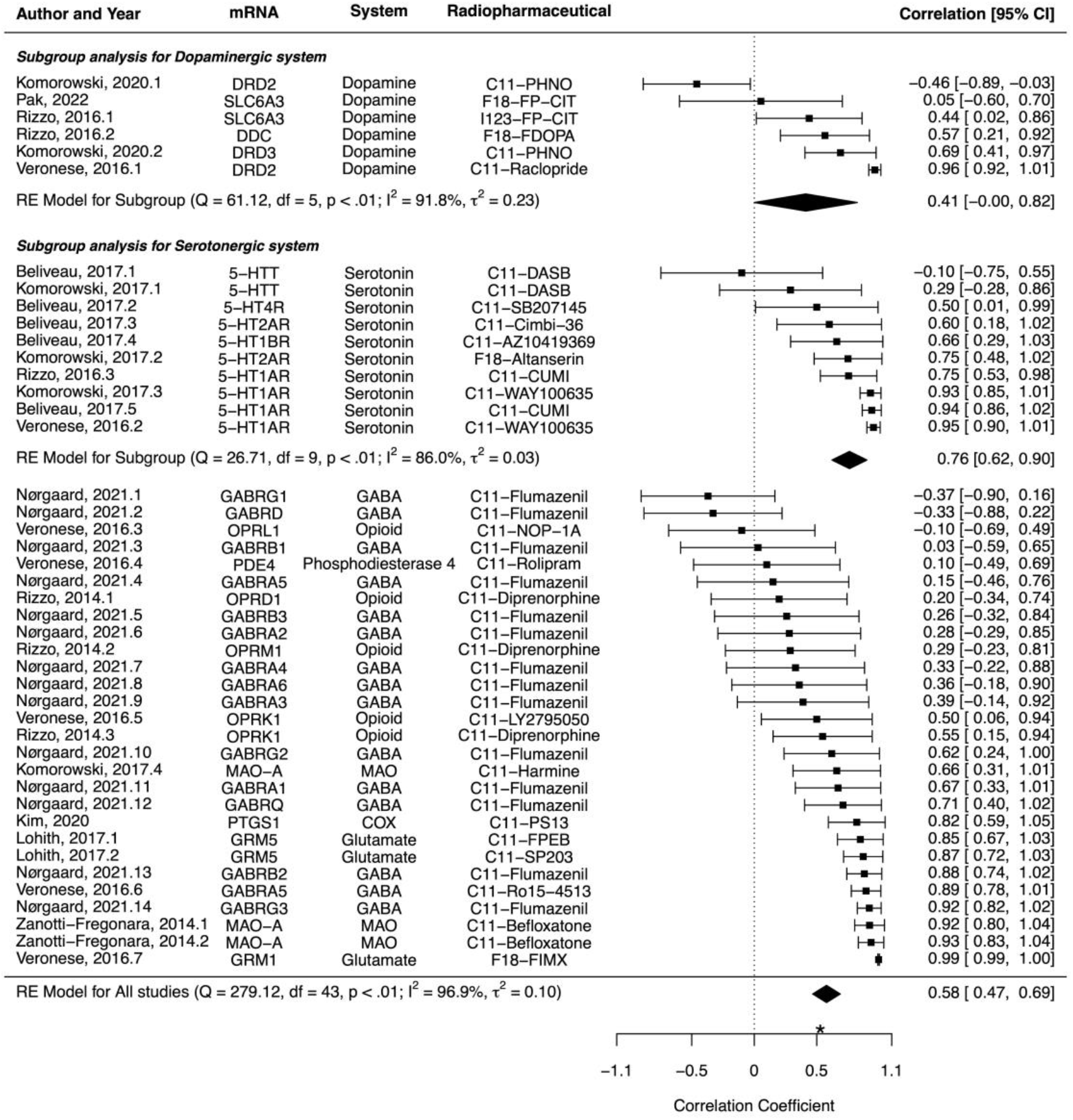
Forest plots for the correlation coefficient between protein expression from PET/single photon emission computed tomography (SPECT) and mRNA expression from the Allen Human Brain Atlas. The pooled correlation coefficient was 0.58 (95% CI: 0.47 ~ 0.69, I^2^=96.9%). In a subgroup analysis, the pooled correlation coefficients were 0.41 (0 ~ 0.82, I^2^=91.8%) for dopaminergic system, and 0.76 (0.62 ~ 0.90, I^2^=86.0%) for serotonergic system, without a significant difference (p=0.1161). *Asterisk indicates the correlation coefficient between CNR1 mRNA expression and CB1 receptor PET in this study.

**Table 1.**
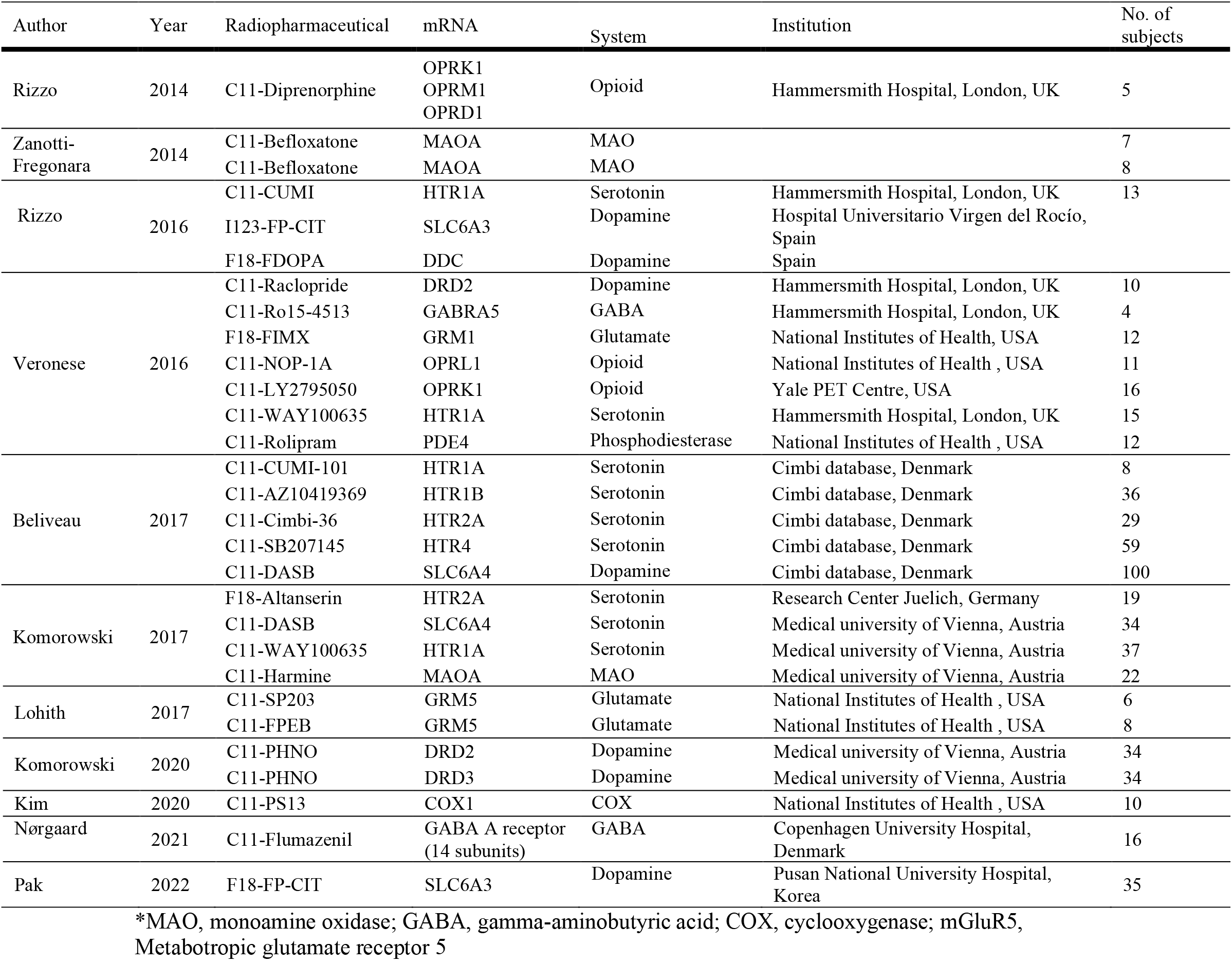
Studies included in meta-analysis.

## 4. DISCUSSION

Our main finding was that CNR1 mRNA expression was moderate to strongly correlated with CB1 receptor availability from V_T_ of F18-FMPEP-d2 PET, showing the predictive power of CNR1 mRNA expression on CB1 receptor. From the meta-analysis, the moderate to strong correlation was observed between mRNA expression from the Allen Human Brain Atlas and protein expressions from PET/SPECT scans across multiple genes, with the pooled correlation coefficient of 0.58, which was similar with the correlation coefficient of 0.5026 between F18-FMPEP-d2 PET scans and CNR1 mRNA in this study.

The Allen Human Brain Atlas is a multimodal atlas of gene expression and anatomy of all brain regions from 6 postmortem brains with microarray-based expression, in situ hybridization gene expression, and magnetic resonance imaging-based brain mapping (Shen et al., 2012). As the Allen Human Brain Atlas provides spatial expression profiles of multiple genes, the correlation study directly shows the predictive power of genes for in vivo protein levels from PET scans. In this regard, to understand brain function of CB1 receptor at the level of both mRNA and protein, the integration of brain mRNA atlas with PET scans and autoradiography were done in this study. These data show that the CB1 expression estimates derived from the radioligand F18-FMPEP-d2 are well in line with those from the ex vivo samples from the Allen Human Brain atlas, providing further evidence for the validity of this radioligand for mapping the endocannabinoid system.

Recently, there has been a growing interest in medical use of cannabinoids. However, there is limited evidence on the use of cannabinoids for treating the mental disorders, as tetrahydrocannabinol worsened the symptoms of psychosis, and increased the adverse events from treatment, and still, few randomized controlled trials examined the effect of medical use of cannabinoids (Black et al., 2019). Actually, adolescents and those with psychotic disorders might be vulnerable to the use of cannabinoids (Hill et al., 2021). The type 1 cannabinoid receptor (CB1) is a key component of the endocannabinoid system, which consists of cannabinoid receptors, endogenous ligands and their metabolic enzymes (Tao et al., 2020). CB1 receptor, encoded by CNR1 gene, is expressed in cortex, hippocampus, amygdala, basal ganglia outflow tracts, and cerebellum (Fig 1 & 2), and these circuits may be responsible for the behavioral effects of cannabis (Mackie, 2005). CB1 receptors are found primarily in the presynapses of the neurons, unlike other receptors of neurotransmitters which are located in the postsynapses (Mechoulam and Parker, 2013). Activation of CB1 receptors leads to a decrease in cyclic adenosine monophosphate accumulation (cAMP), inhibition of cAMP-dependent protein kinase and stimulation of mitogen-activated protein kinase activity. CB1 receptor level is increased from adolescence to adulthood in rats, showing its importance in neurogenesis (Aguado et al., 2006; Verdurand et al., 2011). Also, changes in CB1 receptor have been reported in patients with neuropsychiatric disorders (Basavarajappa, 2007). Therefore, CB1 receptor has become a target for drug development and in vivo imaging biomarker for neuropsychiatric disorders (Van Laere, 2007).

The association between CNR1 mRNA expression and protein expression from either PET scans or autoradiography showed the moderate to strong correlation, despite of samples stemming from unrelated populations. However, we also have to consider that there are many complex and various post-transcriptional mechanisms that are involved in turning mRNA into protein (Rizzo et al., 2016). mRNA transcripts interact with intra, extracellular stimuli, and are modified, regulated by non-coding RNAs (Di Liegro et al., 2014), which have an influence on protein expression for each cell type (Cheng et al., 2005; Rizzo et al., 2014). In addition, technologies regarding measurement of either mRNA or protein expression may not be perfectly accurate (Veronese et al., 2016). Calculation of mRNA expression for each probe has its advantages and disadvantages (Arnatkeviciute et al., 2019). Therefore, probe selection has an impact on the final results of mRNA expression (Arnatkeviciute et al., 2019).

In this study, we selected the median gene expression within each ROI to minimize the possible bias from probes. Also, mRNA expression is analyzed in the cytoplasm, while CB1 receptor is predominantly expressed in the cell membrane, presynaptically (Mechoulam and Parker, 2013). However, genomic atlas does not provide an accurate mapping at a cellular level, and mRNA expression as well as protein expressions from PET scans and autoradiography are averaged within each ROI, yielding representative expression values. Also, autoradiography provides far less spatial information than other technologies (Beliveau et al., 2017).

Previously, the predictive power of brain mRNA mappings of the Allen Human Brain Atlas has been investigated for several PET-derived protein expressions, including serotonin receptors (Beliveau et al., 2017; Komorowski et al., 2017; Rizzo et al., 2014), serotonin transporters (Beliveau et al., 2017; Komorowski et al., 2017), opioid receptors (Rizzo et al., 2014), dopamine receptors (Komorowski et al., 2020) and monoamine oxidase A (MAO-A) (Komorowski et al., 2017; Zanotti-Fregonara et al., 2014). The correlation coefficient ranged from −0.46 for dopamine receptor to 0.99 for glutamate receptor (Figure 4), while the mean correlation across probes and radioligands was 0.58. The presently observed correlations for CB1 *(rho* = 0.5026 and 0.6727) align well with this overall effect. The most commonly analyzed protein, serotonin 1A receptor was investigated with multiple radiopharmaceuticals including C11-CUMI-101 (Beliveau et al., 2017; Rizzo et al., 2016), and C11-WAY100635 (Komorowski et al., 2017; Veronese et al., 2016). Even with the same gene expression of HTR1A mRNA from the Allen Human Brain Atlas, a wide range of correlation coefficients with protein expression has been shown; 0.75 – 0.95, probably due to the radiopharmaceuticals and ROIs included in each study. In addition, the majority of the studies included only a small number of subjects, typically less than 30. However, the moderate to strong correlation was observed between mRNA expression and protein expressions across multiple genes, showing the predictive power of genes to estimate protein levels of human brains. Further studies are needed to examine the association between protein expression based on multiple PET radioligands and gene expression with a uniform method.

## 5. CONCLUSIONS

We observed the moderate to strong associations between gene and protein expression for CB1 receptor in the human brain. Therefore, CNR1 mRNA mapping might have the predictive power for in vivo CB1 receptor protein expression, which was also validated by autoradiography. There have been multiple studies investigating the association between mRNA and protein expression in the human brain, showing the heterogeneous results, however, the moderate to strong correlation was observed between mRNA expression and protein expressions across multiple genes, showing the predictive power of genes to estimate protein levels of human brains. Further study is needed to investigate the relationship between multiple genes and in vivo proteins to improve and accelerate drug development.

## DATA AVAILABILITY

The datasets generated during and/or analysed during the current study are available from the corresponding author on reasonable request.

